# Isolation and characterization of a novel *Wolbachia* bacteriophage from *Allonemobius socius* crickets in Missouri

**DOI:** 10.1101/2021.03.31.437854

**Authors:** Jonah Kupritz, John Martin, Kerstin Fischer, Kurt C Curtis, Joseph R Fauver, Yuefang Huang, Young-Jun Choi, Wandy L Beatty, Makedonka Mitreva, Peter U Fischer

## Abstract

*Wolbachia* are endosymbionts of numerous arthropod and some nematode species, are important for their development and if present can cause distinct phenotypes of their hosts. Prophage DNA has been frequently detected in *Wolbachia*, but particles of *Wolbachia* bacteriophages (phage WO) have been only occasionally isolated. Here, we report the characterization and isolation of a phage WO of the southern ground cricket, *Allonemobius socius*, and provided the first whole-genome sequence of phage WO from this arthropod family outside of Asia. We screened *A. socius* abdomen DNA extracts from a cricket population in eastern Missouri by quantitative PCR for *Wolbachia* surface protein and phage WO capsid protein and found a prevalence of 55% and 50%, respectively, with many crickets positive for both. Immunohistochemistry using antibodies against *Wolbachia* surface protein showed many *Wolbachia* clusters in the reproductive system of female crickets. Whole-genome sequencing using Oxford Nanopore MinION and Illumina technology allowed for the assembly of a high-quality, 55 kb phage genome containing 63 open reading frames (ORF) encoding for phage WO structural proteins and host lysis and transcriptional manipulation. Taxonomically important regions of the assembled phage genome were validated by Sanger sequencing of PCR amplicons. Analysis of the nucleotides sequences of the ORFs encoding the large terminase subunit (ORF2) and minor capsid (ORF7) frequently used for phage WO phylogenetics showed highest homology to phage WOKue of the Mediterranean flour moth *Ephestia kuehniella* (94.18% identity) and WOLig of the coronet moth, *Craniophora ligustri* (96.86% identity), respectively. Transmission electron microscopy examination of cricket ovaries showed a high density of phage particles within *Wolbachia* cells. Isolation of phage WO revealed particles characterized by 40-62 nm diameter heads and up to 190 nm long tails. This study provides the first detailed description and genomic characterization of phage WO from North America that is easily accessible in a widely distributed cricket species.

## Introduction

It is estimated that 66% of all insect species and the majority of filarial parasites that infect humans are infected/colonized with *Wolbachia* [1]. *Wolbachia* causes phenotypes such as cytoplasmic incompatibility and feminization in arthropods, or support growth and reproduction in filarial nematodes [2, 3]. *Wolbachia* is divided into several supergroups based on its ftsZ gene sequence, with supergroups A and B found exclusively in arthropods and supergroups C and D found exclusively in nematodes [4]. Active bacteriophages infecting *Wolbachia* (phage WO) were first discovered in the year 2000 and remain one of few published cases of bacteriophages that infect intracellular bacteria [5]. The persistence of the phage despite its documented lytic activity has led to the hypothesis that phage WO provides benefit to its *Wolbachia* or arthropod host [6]. Phage WO may regulate *Wolbachia* density and therefore, affect development and phenotype of its eukaryotic host [7]. Further, phage WO may supply *Wolbachia* with accessory genes for cytoplasmic compatibility and male killing [8].

In recent years, an increasing number of *Wolbachia* genomes have been sequenced and phage WO is of interest for being the only known mobile genetic element in *Wolbachia*, which is highly resistant to current genetic modification tools, and its hypothesized role in generating the high level of diversity seen among *Wolbachia* today [6, 9]. Evidence has been provided for horizontal gene transfer between *Wolbachia* strains mediated by phages WO [10]. Phages are estimated to infect most of the *Wolbachia* taxa in the supergroups A and B. However, for a majority of these phages, sequence data is limited to the minor capsid protein-coding gene, and there remain entire families and genera of *Wolbachia*-harboring arthropods in which phage has not yet been described [5]. One such example is found in crickets (*Gryllidae*) of the genus *Allonemobius* (ground crickets), whose members include *A. socius* (the southern ground cricket) and *A. maculatus* (the spotted ground cricket), found throughout North America. *Wolbachia* belonging to supergroup B has been identified in *A. socius* (*w*Soc), where it is hypothesized to play a role in altering the length of female crickets’ spermathecal duct [11, 12]. However, phage WO has neither been identified nor described in *Allonemobius*.

In the present study we identified, for the first time, a phage WO in *Allonemobius* crickets (phage WOSoc) and estimated its prevalence. We characterized the novel phage WOSoc by immunohistochemistry, transmission electron microscopy, and whole genome sequencing, expanding the limited set of fully described bacteriophages of *Wolbachia* by adding this novel bacteriophage for which we provide evidence of complete phage particle production, host lysis, and genetic manipulation.

## Materials and Methods

### Sample collection and DNA extraction

Adult *A. socius* crickets (n= 40) were collected in the summer of 2019 from Forest Park, St. Louis, Missouri, USA (N 38.4° 38’, W 90° 17’). Crickets were sexed based on the presence (female) or absence (male) of an ovipositor and ecological data including morphological features and geographical distribution were used to confirm species identification. All insects were euthanized by placement at -20° C for 30 minutes before dissection and homogenization of abdomens in 500 µL of phosphate buffered-saline by 15-minute high-intensity beating with a 3.2 mm chrome Disruption Bead (BioSpec Products, Bartlesville, USA) on the Vortex-Genie 2 mixer (Scientific Industries, Inc., Bohemia, USA). The homogenate was spun down, and DNA was prepared from the supernatant using the DNeasy Blood & Tissue Kit (Qiagen, Hilden, Germany) according to manufacturer recommendations, with elution into 100 µL sterile water and storage at -20°C or 4°C until use.

### PCR for phage and *Wolbachia* detection

Conventional PCR reactions with total cricket abdomen genomic DNA template were run using previously validated primers to the conserved *Wolbachia* surface protein (WSP) gene [13] and to the *Wolbachia* phage capsid protein (WPCP) gene [14]. PCR was performed in 25 µL reactions with 0.625 µL of 10 µM forward and reverse primers (250 nm final concentration), 2 µL DNA template (2-5 ng), 12.5 µL Hot Start Taq DNA Polymerase (2X (New England Biolabs, Ipswich, USA), and 9.25 µL sterile water. Following an initial 30 s denaturation at 95°C, 40 cycles were run with 30 s denaturation at 95°C, 60 s annealing at 55°C, 1 min extension at 68°C, and a single 5 min final extension at 68°C. For each primer set and reaction, sterile water was run as a non-template control. PCR products were sent to Genwiz (South Plainfield, USA) for Sanger sequencing. Forward and reverse primer sequencing reactions were performed for each region of interest and chromatograms were visually inspected for base call quality.

### Real-time PCR prevalence estimates

Primer 3 software [15] was used to create qPCR-optimized WSP and WPCP primers from their respective *w*Soc and WOSoc sequences (Table 1). For each DNA template and primer set, qPCR reactions were performed in duplicate 25 µL reactions with 0.625 µL of 10 µM forward and reverse primers (250 nm final concentration), 2 µL DNA template, 12.5 µL Power SYBR Green Master Mix (Thermo Fisher, Waltham, USA), and 9.25 µL sterile water using the standard Power SYBR Green PCR Master Mix RT-PCR Protocol (Protocol Number 436721) on a QuantStudio 6 Flex Real-Time PCR System (Thermo Fisher). As positive controls for the WSP and WPCP primer sets, we used 2µL Sanger-confirmed WSP-and WPCP-positive cricket genomic DNA. Sterile water was run as the negative control. A conservative cycle threshold (CT) cutoff value of ≤ 23 for positive determination was set for both primer sets based on melting curve and relative abundance analysis corresponding to three standard deviations below the negative control detection level.

**Table 1.**
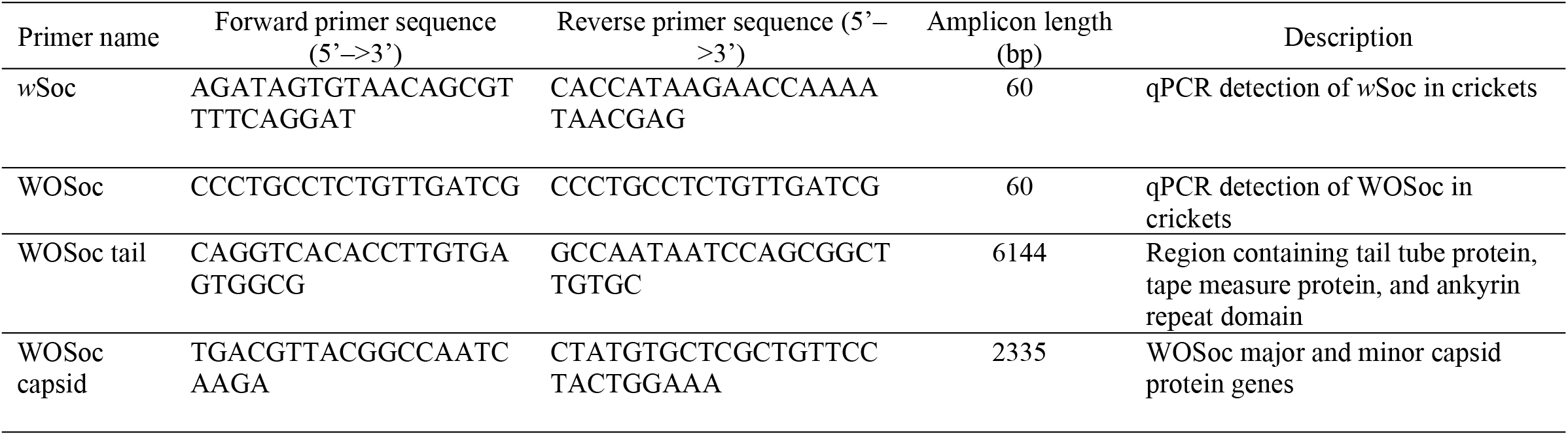
List of primers designed and used in the study.

**Table 2.** Prevalence estimates of *Wolbachia* surface protein (WSP) and phage capsid protein (WPCP) DNA in *Allonemobius socius* crickets from Missouri.

|  |  | WSP |  | Total N (%) |
| --- | --- | --- | --- | --- |
|  |  | Positive N (%) | Negative N (%) |  |
| WPCP | Positive N (%) | 19 (47.5%) | 1 (2.5%) | 20 (50%) |
|  | Negative N (%) | 3 (7.5%) | 17 (42.5%) | 20 (50%) |
|  | Total N (%) | 22 (55%) | 18 (45%) | 40 (100%) |

Estimates are based on a SYBR qPCR assay with a strict cutoff of $CT \leq 23$ in 40 adult *A. socius* abdomen genomic DNA extracts.

### Immunohistology for visualization of *Wolbachia*

For immunohistology, 10 whole *Allonemobius* crickets were fixed in 80% ethanol, embedded in paraffin, and sectioned at 10 µm. Sections were stained with a monoclonal mouse antibody against the *Brugia malayi Wolbachia* surface protein (1:100) for 1 hour at room temperature or overnight at 4°C using the alkaline phosphatase-anti-alkaline-phosphatase (APAAP) technique according to the manufacturer’s protocol (Dako, Carpinteria, CA, USA). All antibodies were diluted in TBS with 0.1% BSA. TBS with 1% albumin was used as a negative control, whereas sections from *B. malayi* worms from previous studies [16] were used as positive controls, respectively. After a 30 min incubation with the secondary rabbit-anti mouse IgG antibody (1:25) (Dako) followed a 30 min incubation step with alkaline-phosphatase-anti-alkaline-phosphatase (1:40) (Millipore Sigma, St. Louis, USA). As substrate, SIGMA*FAST* Fast Red TR/Naphthol AS-MX (Millipore Sigma) Tablets were used, and sections were counterstained with Mayer’s hematoxylin (Millipore Sigma). Sections were analyzed using an Olympus-BX40 microscope and photographed with an Olympus DP70 camera.

### DNA extraction, library preparation and whole genome sequencing

High molecular weight (HMW) DNA was purified from a homogenate of a whole single adult female cricket prepared by 15 min beating with a lead bead using the MagAttract HMW DNA Kit (Qiagen) according to manufacturer specifications, eluting in 100 µL sterile water. Presence of HMW was confirmed by gel electrophoresis. Presence of WPCP in HMW DNA was confirmed by qPCR. DNA was then purified further using AMPure XP beads (Beckman Coulter, Brea, USA) at a ratio of 1.8:1 bead to DNA sample. Library was prepared according to Oxford Nanopore’s 1D Genomic DNA Ligation Protocol (Version GDE_9063_v109_revA) using the LSK-109 Ligation Sequencing Kit (Oxford Nanopore Technologies, Cambridge, England) with DNA fragments of all sizes purified using the Short Fragment Buffer. 60 µL of library containing 12 µL genomic DNA was loaded as input into the flow cell and the sequencing reaction run for 20 hours using MinKNOW GUI software (Oxford Nanopore Technologies) set to the High Accuracy Flip-Flop Model, generating 6.1 giga base pairs of data. Basecalling of Fast5 files into Fastq format was performed using Guppy neural network basecalling software [17]. Base statistics, average quality per read, sequence duplication level, and GC content were assessed using FastQC software (Babraham Institute, Cambridge, UK). In parallel, genomic DNA was extracted from the ovary tissue of a single cricket using Qiagen DNeasy kits as described above and sequenced using a NovaSeq 6000 Sequencing System (Illumina, San Diego, USA) with 2×150 bp output generating 12.2 giga base pairs of data, following qPCR confirmation of phage positivity in the sample

### Assembly and annotation of the WOSoc genome

Putative WOSoc reads were extracted by mapping MinION sequences against published phage WO reference genomes using Minimap2 software [18]. Mapped reads were then mapped against themselves in order to merge overlapping reads. The self-mapping output and the MinION-generated Fastq sequences were input into CANU Single Molecule Sequence Assembler [19] to generate a phage assembly consisting of multiple contigs. Quality trimming and adapter clipping of Illumina reads was performed using Trimmomatic [20]. The PRICE assembly tool [21] was used to extend existing contigs using the Illumina data. Redundans was used collapse redundant contigs, scaffold contigs, and close gaps using both the Oxford Nanopore Technologies (ONT) reads and Illumina reads. ONT reads were error-corrected using FMLRC [22] before feeding them into the Redundans pipeline [23]. We then manually curated the assembly and corrected assembly errors. Finally, Pilon automated genome assembly improvement pipeline [24] was used to polish the assembly and reduce base-call errors. Annotation of the assembled phage genome was performed using the Rapid Annotation Using Subsystem Technology Toolkit (RASTtk) SEED-based prokaryotic genome annotation engine with default presets, which has established validity for annotating phage genomes [25, 26], identifying genomic “features” (protein-coding genes and RNA). Genomic features were visualized in scaffolds independently and manually color-coded by function using Gene Graphics visualization application [27].

### PCR and Sanger sequencing for genome verification

Primers were manually designed to amplify phage tail and capsid regions based on the MinION reads (Table 1). Conventional PCR reactions were run with these primers and cricket abdomen DNA as described previously with a 60°C annealing temperature for both primer sets. Amplicons were gel-excised, purified, and 3730 Sanger sequenced.

### Phylogenetic analyses

DNA sequences of phage WO open reading frames 2 (ORF2) and 7 (ORF7), respectively coding for the large terminase subunit and minor capsid, are biomarkers known to produce highly congruent phage WO phylogenies [5]. Nucleotide sequences of ORF2 and ORF7 of WOSoc were compared to published gene sequences in NCBI Genbank. Phylogenetic trees were generated based on WOSoc ORF2 and ORF7 identity to the top 4 BLAST hits based on pairwise alignments using the NCBI BLAST Tree View Neighbor-Joining tree method with distances from the node computed by NCBI presets. ORF2 sequence was extracted from Scaffold 1 of the phage assembly, while the entire ORF7 gene was provided by Sanger sequencing of the capsid region as described above.

### Phage particle purification

Phage was purified according to the protocol described in [28] with slight modification. Unless otherwise noted, all reagents were purchased from Sigma-Aldrich, St. Louis, USA. Complete mature *A. socius* males and females (N = 70) were euthanized and thoroughly homogenized in 40 mL of SM buffer (50 mM Tris-HCL, pH 7.5, 0.1 M NaCl, 10 mM MgSO_4_ • 7 H_2_O and 0.1% w/v gelatin containing 1 µg/mL RNase A). Homogenate was incubated on ice for 1 hour followed by 11,000xg centrifugation for 10 minutes at 4°C to remove debris. Solid polyethylene glycol (PEG) was added to homogenate to a final concentration of 10% and mixed by manual shaking for 1 minute, followed by an additional 1-hour incubation on ice and 11,000xg centrifugation for 10 minutes at 4°C. Supernatant was discarded and the remaining pellet was resuspended in 10 mL of SM buffer. To the suspension, an equal volume of chloroform was added followed by centrifugation at 3,000xg for 15 minutes at 4°C to remove the PEG. The aqueous layer containing phage was filtered through a 0.22 µM vacuum filter to remove *Wolbachia* and other bacteria. Phage lysate was concentrated using Amicon Ultra-15 100 kDA Centrifugal Units (Millipore, Burlington, USA) according to [29] and reconstituted in a final volume of 1 mL of SM buffer.

### Transmission electron microscopy (TEM) for visualization of WOSoc particles

From freshly caught adult female *A. socius*, ovaries were dissected and adsorbed to an electron transparent sample support (EM) grid. Tissue was washed in PBS and fixed in 1% glutaraldehyde for 5 minutes at room temperature, followed by two 30-second washes with deionized water. Phage particles were negatively stained in 1% uric acid for 1 minute and wicked gently and placed in a grid box to dry. Phage suspension was processed identically, with 50 µL of the concentrated suspension adsorbed to an EM grid. Samples were observed on a JEOL 1200 EX transmission electron microscope (JEOL USA Inc., Peabody, USA) equipped with an AMT 8-megapixel digital camera (Advanced Microscopy Techniques, Woburn, USA) To confirm the presence of phage in *Wolbachia* by TEM, one half of the ovaries of each of 6 crickets was fixed in 2% paraformaldehyde/2.5% glutaraldehyde (Polysciences Inc., Warrington, USA) in 100 mM phosphate buffer, pH 7.2, for 1 hour at room temperature. The other half of the ovary sample was added to 1X PBS for DNA extraction and confirmation of *Wolbachia* presence by PCR. Only samples that were positive by PCR for Wolbachia were further processed for TEM. These samples were washed in phosphate buffer and post-fixed in 1% osmium tetroxide (Polysciences Inc.) for 1 hour. Samples were then rinsed extensively in distilled water prior to staining with 1% aqueous uranyl acetate (Ted Pella Inc., Redding, USA) for 1 hour. Following several rinses in distilled water, samples were dehydrated in a graded series of ethanol and embedded in Eponate 12 resin (Ted Pella Inc.). Sections of 95 nm were cut with a Leica Ultracut UCT ultramicrotome (Leica Microsystems Inc., Bannockburn, USA), stained with uranyl acetate and lead citrate, and viewed on a JEOL 1200 EX transmission electron microscope (JEOL USA Inc.) equipped with an AMT 8-megapixel digital camera (Advanced Microscopy Techniques) [30].

## Results

### Prevalence of Phage WO and *Wolbachia* in *A. socius*

DNA encoding WSP was used as a marker for assessing the prevalence of *Wolbachia* in crickets. In order to confirm the DNA sequence of WSP of Missouri crickets, DNA was amplified by conventional PCR using the pre-validated WSP primers. WSP sequence showed 100% identity to WSP of *A. socius* from Virginia (Accession: AY705236.1, data not shown). A 400 bp amplicon of phage DNA was amplified by conventional PCR using pre-validated primers corresponding to nucleotide positions 7353-7761 of phage WO of cricket *Teleogryllys taiwanemma* cricket and showed close homology to the capsid protein genes from phage WO of *Supella longipalpa* (95.50% identity, 100% query coverage, Accession: KR911861.1) and *Cadra cautella* (94.50% identity, 100% query coverage, Accession: AB478515.1). The *A. socius* WSP and phage WOSoc WPCP gene sequences were used to design SYBR-based real-time PCR assays for WSP and WPCP, respectively. Using the strict CT cutoff of 23 cycles, we determined that from 40 insects sampled 19 (47.5%) were positive for both WPCP and WSP DNA via qPCR with our optimized primers; three samples (7.5%) were WSP-positive but WPCP-negative.

Confirmation of the *Wolbachia* prevalence results was done using an orthogonal approach, i.e visualization by immunohistology. Endobacteria were found in about 50% of the female crickets. They were detected throughout the abdomen, however density was highest in the reproductive tract (Fig. 1). *Wolbachia* were detected in distinct, but varying parts of the panoistic ovarioles. In the apical part of the ovariole, *Wolbachia* were seen in the inner section of the follicle epithelium (Fig. 1C), but in more mature eggs, these cells are devoid of *Wolbachia* and endobacteria were concentrated in large numbers in one pole of the egg cell (Fig. 1F). The high density of *Wolbachia* in developing eggs ensures transovarial transmission of *Wolbachia* and phage WO [31]. It is expected that in this context, where *Wolbachia* negatively impacts its host’s fitness, host selection will act to limit or eliminate the endosymbiont, which may explain the less than ubiquitous *w*Soc prevalence. At the same time, high phage density favors the insect host in a parasitic *Wolbachia* context, which benefits from the reduction in *Wolbachia* density resulting from phage-mediated lysis or transcriptional regulation, which could promote phage abundance to the high levels seen in *w*Soc-infected insects [6].

**Figure 1.**
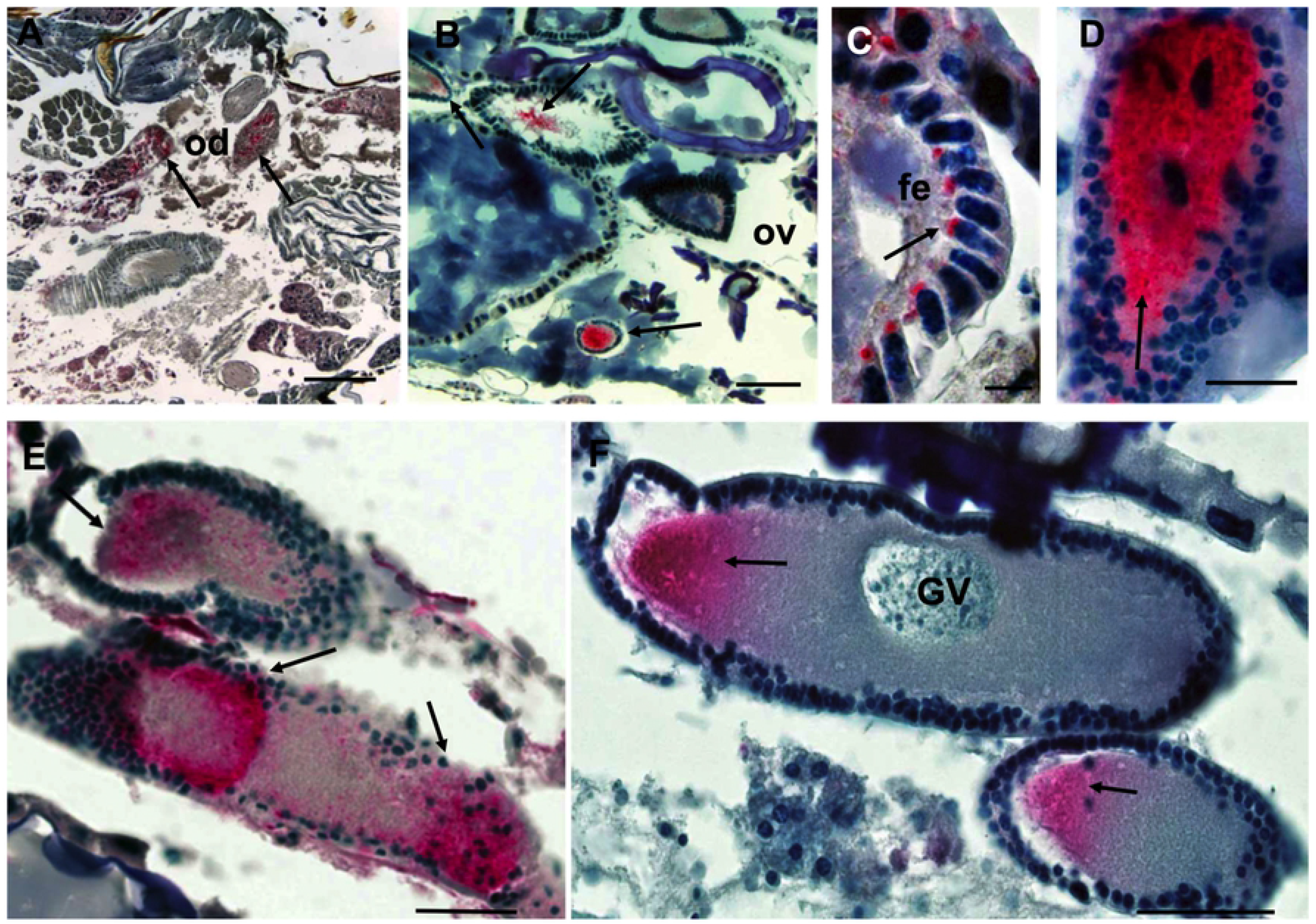
Immunohistological localization of *w*Soc. Black arrows indicate *Wolbachia* (red). **A**. Posterior abdomen containing intestinal tissue and oviduct containing *Wolbachia* (200μm). **B**. Ovary tissue showing dense clusters of *Wolbachia* at the site of maturing oocytes (200μm). **C**. *Wolbachia* localized to the follicle epithelium. **D** (50 μm), **E**, and **F**. Close-up of oocytes in the female cricket oviduct showing *Wolbachia* cells in studding follicles. The nucleus (GV) is visible in the upper oocyte in **F. (**20 μm) Abbreviations: FE = follicle epithelium; od = oviduct; ov = ovaries; GV = germinal vesicle. Scale bar: 10 µm.

### Isolation and visualization of Phage WO of *A. socius*

Although we detected capsid DNA of phage WO in most *Wolbachia*-positive *A. socius* samples, it was theoretically possible that this was exclusively prophage DNA incorporated into the genome of *Wolbachia* and that no phage particles were formed. Therefore, we used TEM to visualize particles of phage WO of *A. socius*. Several intracellular *Wolbachia-*containing stereotypical hexagonal phage particles were detected in ovarian tissue (Fig. 2). Small clusters of *Wolbachia* cells that contained up to 30 complete phage particles per cells were obverted to mature egg cells (Fig 2. A, B, D). TEM examination of the filtrate from phage precipitation revealed numerous phage WO particles. Measurement of 10 particles showed an average diameter of the icosahedral head structure of 47 and 62 nm (±x nm SD) and 175 and 135nm long, striated tails (Fig 2. E, F).

**Figure 2.**
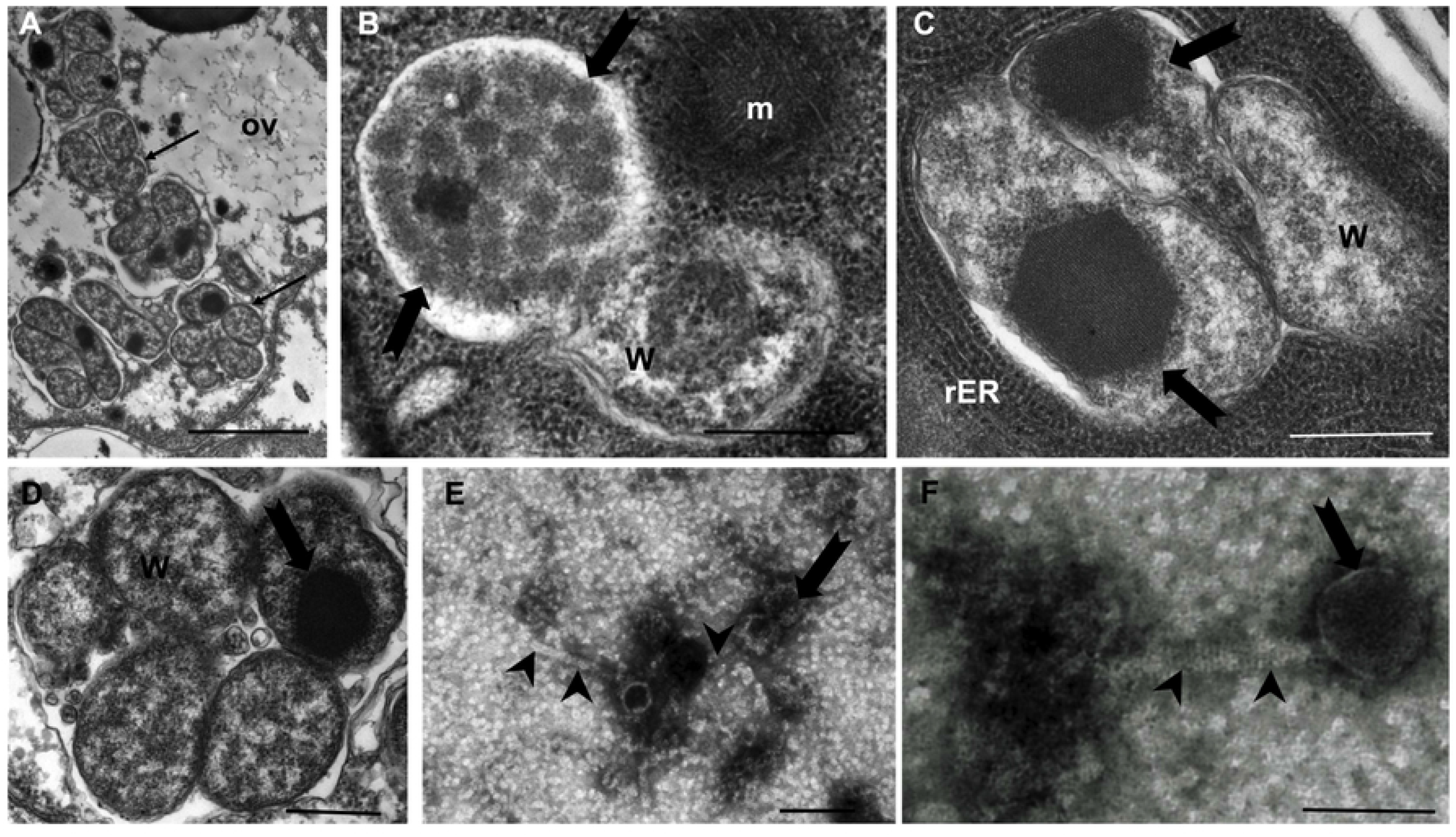
Transmission electron microscopy (TEM) of WOSoc particles. **A**. Clusters of intracellular *Wolbachia w*Soc (arrows) in the ovary of *A. socius* (scale bar 2 μm). **B**. Densely packed phages WOSoc (arrows) inside a *Wolbachia* endobacterium (scale bar 500 nm). **C**. and **D**. Compact, electron dense hexagonal arrays of phages WOSoc (arrows) in *Wolbachia* (scale bar 500 nm). **E**. and **F**. Complete, purified phage particles with 47 to 62 nm capsids (arrow) and 175 to 130 nm tails (arrow head, scale bar 100 nm). Abbreviations: ov, ovaries; W, *Wolbachia*, rER, rough endoplasmic reticulum; m, mitochondrion.

### The WOSoc genome indicates potential for lysis and transcriptional manipulation of the host

Following the detection of phage DNA in WSP-positive crickets and the demonstration of distinct phage particles, we set out to genomically characterize the novel phage WO to gain insight into its lytic potential and its similarity to known phages WO. Using the well-characterized genome of WOVitA1 (a *Wolbachia* bacteriophage found in the parasitic wasp, *Nasonia vitripennis*) as a reference genome, we identified 511 homologous WOSoc reads from the MinION run of whole-cricket homogenate HMW DNA with an average quality per read (Phred Score) of 23, corresponding to an overall base call accuracy exceeding 99%. From these reads, we assembled 12 contigs totaling 53,916 bp at an average depth of 14.6X and a GC content of 35%. After confirming and extending these contigs with Illumina reads and removing low quality reads and reads derived from the *Wolbachia* genome, the WOSoc genome was captured in 4 high-quality scaffolds totaling 55,288 bp. To further validate our assembly, we Sanger sequenced PCR-amplified phage sequence from taxonomically important phage regions using primers generated from the scaffolds, collectively representing nearly one-eighth of the assembly including a continuous, 6,144 bp contig containing complete open reading frames for tail morphogenesis proteins and a 2,289 bp region encoding the major and minor capsid proteins and head decoration protein (all sequence data are available in Supplementary File S1 and the assembly is available in GenBank under the accession IDs MD788653-MW788656). RASTtk annotation identified 63 features which included 33 described and 30 hypothetical or unidentified ORFs based on similarity and bidirectional best hit computation (see Supplementary File S2 for a complete list of these features including full-length protein and gene sequences). Of the 33 described ORFs, over half (N = 17) encoded structural features including tail (N = 9), head (N = 5), and baseplate (N = 3) assembly. We also identified genes necessary for phage replication (N = 2), *Wolbachia* cell wall lysis (N = 3), and a resolvase protein which may catalyze site-specific bacteriophage DNA integration [32] (Fig. 3). Strikingly, we found five features which may regulate *Wolbachia* host transcriptional processes including N-acetylglucosamine-1-phosphate uridyltransferase, which may regulate *Wolbachia* transcription by altering glutamine synthetase activity [33] and glycosyl transferase, which is known to protect phages from bacterial endonucleases [34]. Collectively, these features suggest that WOSoc is an active particle-forming phage with potential for lytic and lysogenic behavior, reflecting an intimate interaction with its bacterial host.

**Figure 3.**
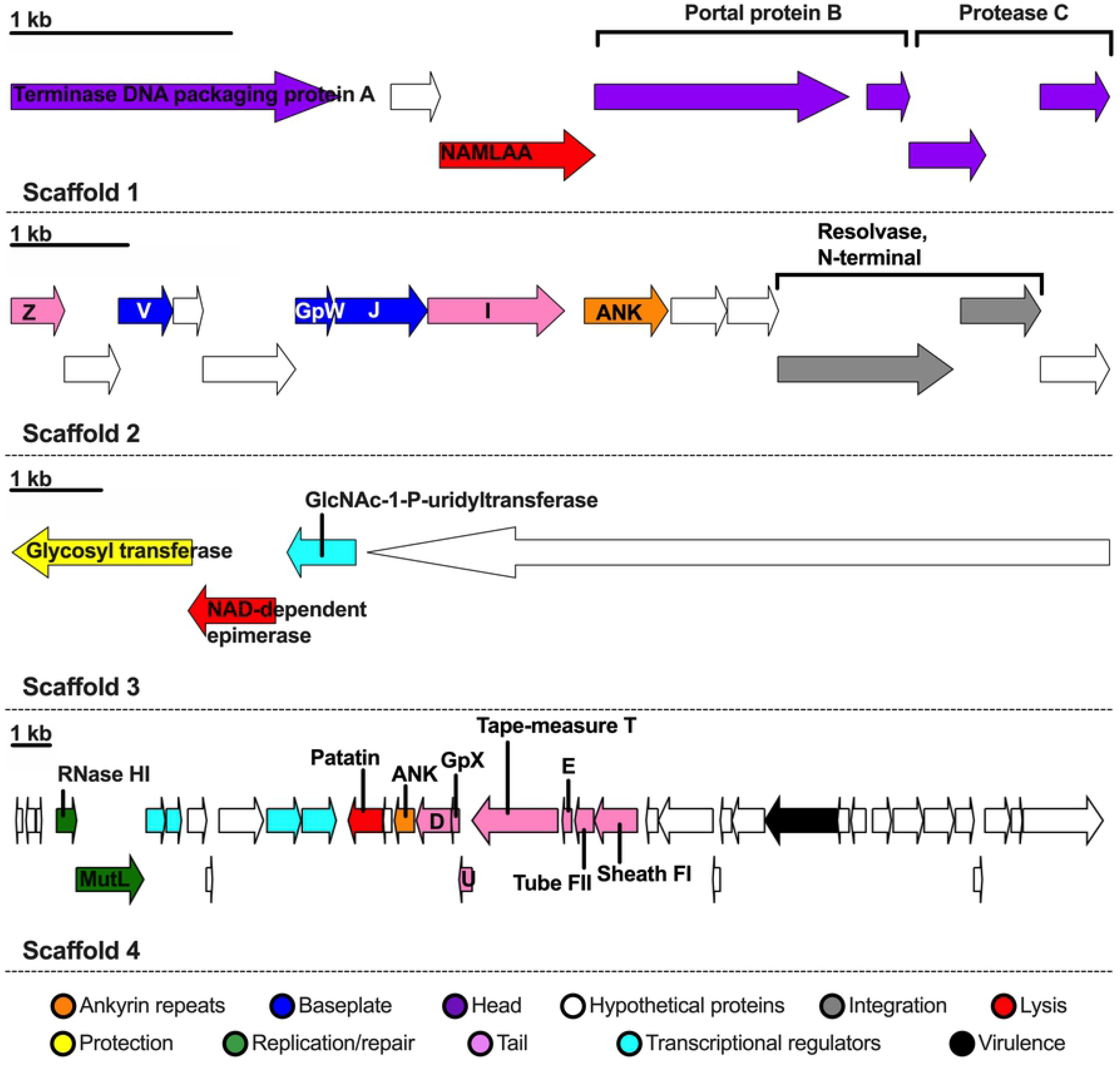
Annotation of the WOSoc genome. 63 features from the RASTk annotation of the 4-scaffold WOSoc assembly are displayed: ankyrin repeats (N = 2), baseplate assembly (N = 3), phage head (N = 5), integration into *Wolbachia*’s genome (N = 2), lysis of *Wolbachia* cells (N = 3), protection from *Wolbachia* endonucleases (N = 1), DNA replication and mismatch repair (N = 2), tail formation (N = 9), transcriptional regulation (N = 5), virulence (N = 1), undescribed hypothetical proteins (N =30). Abbreviations: NAMLAA = N-acetylmuramoyl-L-alanine amidase; ANK = ankyrin. Scale bars: 1 kb within their respective scaffolds.

### Phylogenetic analysis of WOSoc suggests a close relationship with phages WO of moths

In order to compare phage WOSoc to a larger number of phage WO for which the complete genome sequence is not available, we performed pairwise comparison with published ORF2 and ORF7 phage WO sequences. Phage WOSoc ORF2 showed the highest homology to phage WOKue of the Mediterranean flour moth *Ephestia kuehniella* (94.18% nucleotide identity, 100% query coverage, Accession: AB036666.1), while phage WOSoc ORF7 was most similar to WOLig of the coronet moth, *Craniophora ligustri* (96.86% nucleotide identity, 100% query cover, Accession:LR990976.1), both insects of the order Leptidoptera. (Fig. 4). High homology (> 99% identity), which is not uncommon for known conserved phage element sequences, such as the large terminase subunit or the minor capsid protein region, was not observed.

**Figure 4.**
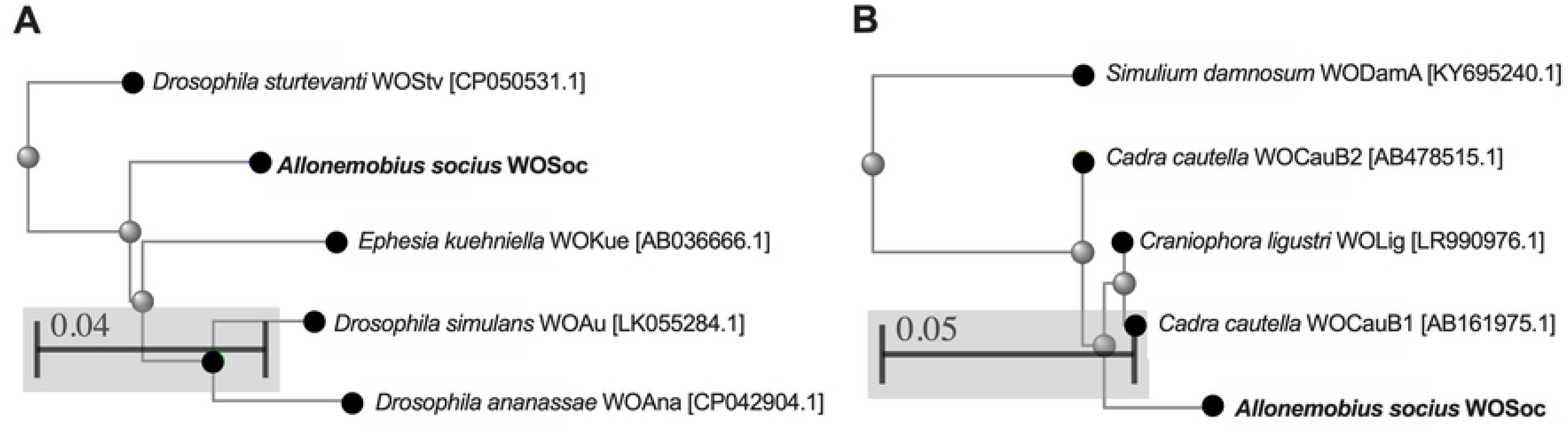
Phylogenetic comparison of WOSoc with published phage sequences. Neighbor-joining trees generated from published phage WO nucleotide sequences aligned to WOSoc **A**. Large terminase subunit (ORF2), showing homology to WOKue of the moth *Ephestia kueniella* and **B**. minor capsid protein (ORF7), showing high homology to WOLig of the moth *Craniophora ligustri*. Scale bars denote distance from the node as calculated by the NCBI Tree View software.

## Discussion

The present study identified for the first time a particle-forming phage WO in North American crickets and provided the whole genome sequence of phage WOSoc. About half of female *A. socius* crickets screened by PCR contained *Wolbachia*. Within arthropod populations, *Wolbachia* infection prevalence closely resembled that seen in other supergroup B infected species [35-38]. More than 85% of *Wolbachia*-positive crickets were also positive for phage WO DNA, indicating co-transmission of *Wolbachia* and phage WO. In a DNA extract of one cricket, we detected phage WO DNA, but not *Wolbachia* DNA. This may be due to contamination with DNA from a phage-positive sample or more likely due to failure of the assay to pick up very low amounts of *Wolbachia* DNA, since a single *Wolbachia* cell may contain many genomes of phage WO. Our TEM examination of *Wolbachia* illustrated this very nicely.

Immunohistological detection of *Wolbachia* in *A. socius* showed high densities of endobacteria in maturing egg cells. TEM examination of ovaries of *A. socius* revealed numerous phage WO particles arranging in varying structures within the *Wolbachia* cells. Occasionally, intracellular, electron-dense, hexagonal arrays where detected that could be the product of phage WOSoc self-assembly into ordered nanoarrays as seen in other bacteriophages [39]. Little information is available that describes the ultrastructure of assembled phage WO particles within *Wolbachia*, however the observed morphology of isolated phage WOSoc particles is similar to other isolated phage WO particles [40-42].

Genomic evidence showed the potential of complete phage WOSoc particle formation and validated the morphology results. Previous reports link the presence of prophage WO DNA with host phenotypes [43, 44]. However, our study showed not only the presence of prophage WO DNA, but also demonstrates particle formation and active propagation of phage WOSoc. Phage particles are the driver of genetic elements into new *Wolbachia* strains. Bacteriophages are considered to be relatively host-specific, but potential host species can be predicted based on sequences of annotated receptor-binding proteins [45]. Unfortunately, these sequences are not always available and further experimental studies have to elucidate the host range of phage WOSoc and its potential to genetically manipulate *Wolbachia*. The isolation of phage WOSoc offers exciting possibilities for understanding the evolutionary and current role of *Wolbachia*’s only known mobile genetic element and an active regulator of *Wolbachia* density on the endosymbiont-induced characteristics such as cytoplasmic incompatibility and reproductive support. Future studies may show whether phage WOSoc plays a role in the spermathecal duct shortening which is a well-documented effect of *Wolbachia* in *Allonemobius* genus crickets [11].

So far, there are only a handful of complete phage WO genome sequences available in the public databases, and this study has expanded the list by adding a validated 55 kilobase genome of phage WOSoc. Like closely related active phage WO of *Cadra cautella*, WOSoc contains intact open reading frames encoding proteins essential to phage particle formation, including tail morphogenesis and DNA packaging, which are absent in inactive, prophages of *Wolbachia* [46].

*Wolbachia* are considered as targets for alternative chemotherapy of human filariasis, caused by parasitic nematodes [47] and as alternative tools for vector control [48]. Therefore, a better understanding of the role of phage WO in regulating *Wolbachia* populations is important to optimize these intervention strategies. In addition, our discovery of a novel phage WO in a common and easily accessible insect species, may help to add *Wolbachia* to the list of bacteria that can be targeted by phage therapy. The concept of phage therapy is old, but has gained new interest in recent years by the rapid increase of antimicrobial resistance [49]. Future studies are needed to show whether phage WOSoc can be utilized to manipulate *Wolbachia* in *A. socius* or other host species infected by *Wolbachia*.

## Supporting Information

S1 Assembly of the genome of phage wAsoc and selected confirmed DNA sequences used for phylogenetic analysis.

S2 Annotation of the genome of phage wAsoc.

## Acknowledgements

The authors would like to thank Dr. Gary Weil for his support. The study was funded in part by The Foundation for Barnes Jewish Hospital. JK was supported by the Washington University Biology Summer Undergraduate Research Fellowship Program (BIOSURF).

